# Machine Learning-Based Enzyme Engineering of PETase for Improved Efficiency in Degrading Non-Biodegradable Plastic

**DOI:** 10.1101/2022.01.11.475766

**Authors:** Arjun Gupta, Sangeeta Agrawal

**Author notes:** **Correspondence:** Sangeeta Agrawal, MD. Arjun Gupta,.

## Abstract

Globally, nearly a million plastic bottles are produced every minute (1). These non-biodegradable plastic products are composed of Polyethylene terephthalate (PET). In 2016, researchers discovered PETase, an enzyme from the bacteria *Ideonella sakaiensis* which breaks down PET and nonbiodegradable plastic. However, PETase has low efficiency at high temperatures. In this project, we optimized the rate of PET degradation by PETase by designing new mutant enzymes which could break down PET much faster than PETase, which is currently the gold standard. We used machine learning (ML) guided directed evolution to modify the PETase enzyme to have a higher optimal temperature (Topt), which would allow the enzyme to degrade PET more efficiently.

First, we trained three machine learning models to predict Topt with high performance, including Logistic Regression, Linear Regression and Random Forest. We then used Random Forest to perform ML-guided directed evolution. Our algorithm generated hundreds of mutants of PETase and screened them using Random Forest to select mutants with the highest Topt, and then used the top mutants as the enzyme being mutated.

After 1000 iterations, we produced a new mutant of PETase with Topt of 71.38°C. We also produced a new mutant enzyme after 29 iterations with Topt of 61.3°C. To ensure these mutant enzymes would remain stable, we predicted their melting temperatures using an external predictor and found the 29-iteration mutant had improved thermostability over PETase.Our research is significant because using our approach and algorithm, scientists can optimize additional enzymes for improved efficiency.

## Introduction

According to the UN, over 200 million tons of plastic are produced every year, and 91 percent of all plastic produced is not recycled (1, 2). One non-polluting recycling and waste management method for plastic is enzymatic recycling. In 2016, researchers in Japan stumbled upon a bacteria that consumed and successfully degraded Polyethylene terephthalate (PET), the most common form of non biodegradable plastic. This bacteria was named *Ideonella sakaiensis*. The bacteria contained the two enzymes, PETase and MHETase. PETase was the enzyme that enables *Ideonella sakaiensis* to successfully degrade PET (3).

However, PETase from *Ideonella sakaiensis* has a low efficiency in breaking down PET. Factors that affect the rate of PET degradation by PETase include surface topology of the enzyme, water absorbency of PET, and higher enzyme reaction temperatures (4). Changing these factors can result in faster rates of PET degradation by PETase. For maximum efficiency, the optimal reaction temperature (Topt) should be above 60-65 degrees Celsius. This is because the polymer chain of the plastic fluctuates at these temperatures. This fluctuation allows water molecules to enter between the chains and weaken them, thus improving the efficiency at which an enzyme can break down PET (4).

Directed evolution is the process of performing natural selection on biological molecules such as enzymes, amino acids, *etc* and generating mutants which are then steered toward a user defined goal. First, directed evolution generates different possible mutations of an enzyme. Second, based on those mutations, corresponding mutants are produced and then scored in the lab.The best scoring mutants are then selected based on the user defined goal such as activity or thermostability. Directed evolution then repeats this process with the top mutants from the previous iteration now acting as the main enzyme. Directed evolution was developed by Frances Arnold and won the Nobel prize in 2018 (5).

Performing directed evolution using machine learning on the computer is known as *in silico* directed evolution. For *in silico* directed evolution, instead of producing those mutants in the lab, machine learning is used to score and evaluate different possible mutations of enzymes. Based on the machine learning scores, the algorithm then selects the best mutant and uses it as a starting point again. Machine learning (ML) guided directed evolution is beneficial because machine learning algorithms can take in more data at once, iterations are faster, and the process is cheaper and less time consuming than actually performing directed evolution in the lab. One challenge that exists with this method is that if the machine learning algorithms written do not have a high performance, directed evolution will not achieve its purpose. Another challenge is that the machine learning models that make up the algorithm often perform better with classification tasks rather than predicting a continuous score. One case where ML-guided directed evolution has been used successfully for enzyme engineering is in 2019, where one group of researchers engineered a new enzyme for stereodivergent carbon–silicon bond formation, a new-to-nature chemical transformation (6). However, machine learning guided directed evolution has not previously been used to engineer enzymes which break down non-biodegradable plastic.

We hypothesize that machine learning can be used to predict an enzyme’s optimal temperature, and machine learning can be combined with directed evolution to engineer an optimized mutant of PETase with a Topt greater than 60°C for more efficient breakdown of PET and nonbiodegradable plastic.

In this project, machine learning was used to perform *in silico* directed evolution on the PETase enzyme to design a mutant which has predicted Topt of 70°C, nearly double that of the wild type PETase. This novel enzyme has the potential to break down non-biodegradable plastic more efficiently and at a faster rate than PETase by functioning at a higher optimal temperature. This enzyme is also predicted by external algorithms to have a higher thermostability than the original PETase enzyme. This approach is novel because it is the first to optimize the PETase enzyme’s optimal reaction temperature using a machine-learning guided directed evolution approach.

First, I used enzymes from the Brenda database to train the Linear Regression, Logistic Regression and Random Forest to predict an enzyme’s Topt with high performance. Then we performed *in silico* directed evolution on PETase by generating mutant enzymes and then screening them against the Machine Learning models to predict the new mutant enzyme’s Topt and select new mutants for further evolution.

## Results

A Machine-learning guided directed evolution algorithm was written in Python in order to engineer PETase for a higher thermostability. A flowchart of the approach is shown in Figures 1, 2 and 3. In order to guide the directed evolution, a machine learning model was written to predict an enzyme’s optimal reaction temperature (Topt). Three machine learning models: Random Forest, Linear Regression and Logistic Regression, were trained for this task using enzymes from the BRENDA database. In the second stage, my algorithm generated millions of mutants of PETase by randomly mutating different positions of the amino acid sequence and scored the mutants using Random Forest Regression – the ML model which performed the best – to determine which mutation would lead to the highest Topt. The algorithm then reenacted this mutation and selection process with the best scoring mutants, which now acted as the starting point for the next round of directed evolution.

**Figure 1:**
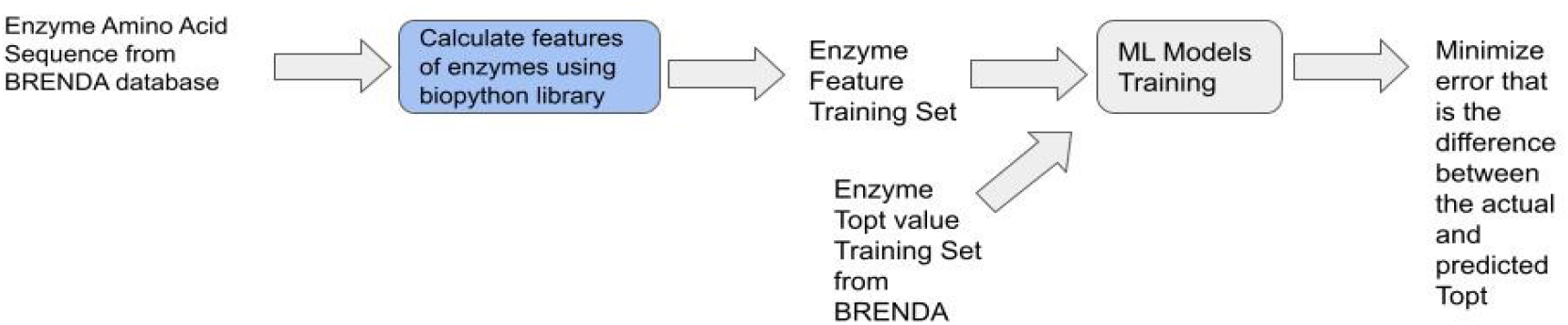
First Flow Chart of Training Stage Approach. This flowchart shows the training of Linear Regression, Logistic regression and Random Forest to predict the Topt of the training set of the data.

**Figure 2:**
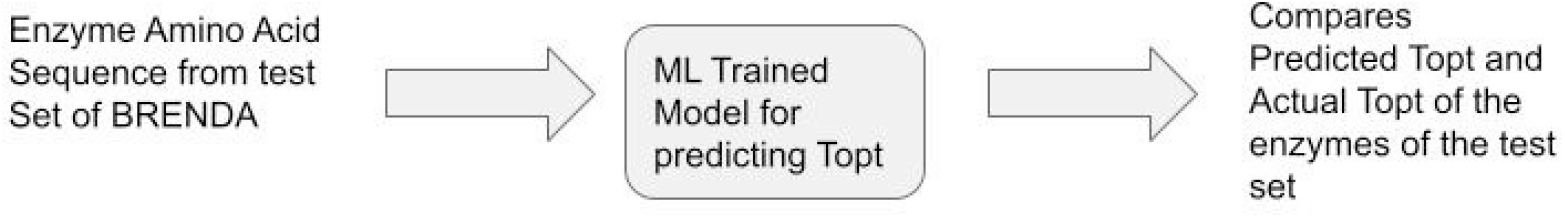
Second Flow Chart of Training Stage Approach. This flowchart shows how the three Machine Learning models were used to predict the Topt of the enzymes in the test set of the data.

**Figure 3:**
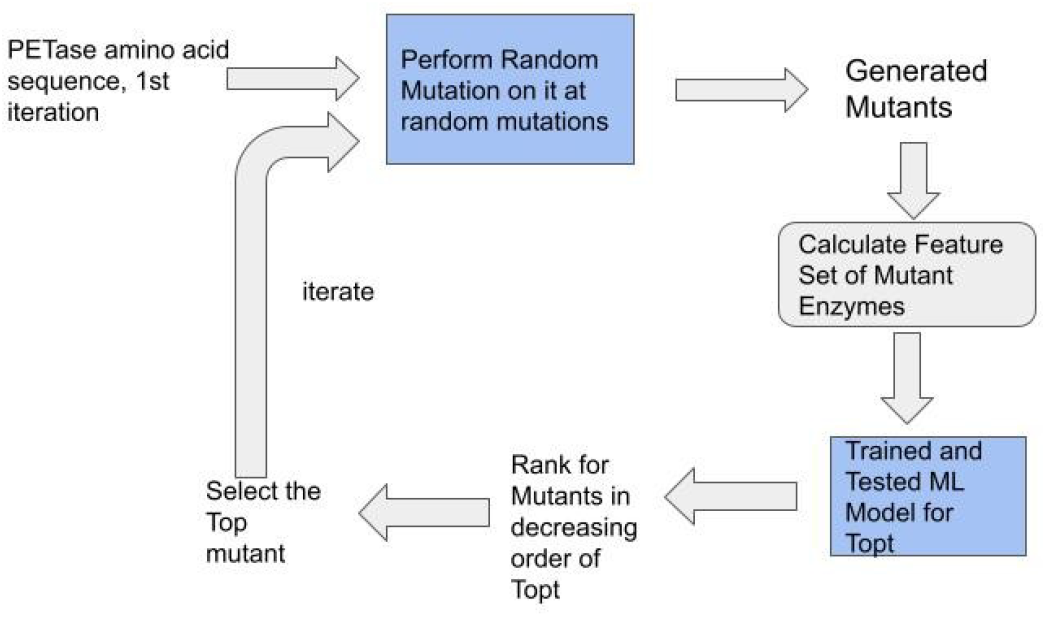
Third Flow Chart show Ml-guided Directed Evolution Approach. This flowchart shows our process of performing in silico directed evolution on PETase.

### Machine-Learning Models for Predicting Optimal Reaction Temperature

Three machine learning models were trained to predict an enzyme’s Topt. Random Forest and Linear Regression were used to predict the actual Topt value of the enzymes. Logistic Regression was used as a classification model to predict whether the enzyme had a Topt above 65 degrees Celsius.

The data set for the machine learning models consisted of enzymes from 11,420 organisms in total obtained from the BRENDA Database (7). Enzymes without Topt values were removed from the dataset. Before performing a cleanup of the data set, there were 2745 enzyme amino acid sequences comprising the dataset. During the cleanup process, we dropped duplicates, amino acid sequences with a length less than or equal to 7 and enzymes with a Topt equal to or lower than 0 degrees Celsius. The final training data consisted of 2643 enzyme amino acid sequences listed with the experimental Topts of each enzyme. The inputs for the models are the enzyme features such as molecular weight, amino acid frequencies, dipeptide frequencies, and the enzyme’s host organism’s optimal growth temperature, as shown in Table 1. These enzyme features were calculated from the amino acid sequence listed from the BRENDA database.

**Table 1.** Enzyme Features calculated for predicting Topt using machine learning models.

| Input Feature | Description | Number of Features |
| --- | --- | --- |
| Global Protein Features | The global protein features included in this list are : Aromaticity, Hydrophobicity, Instability Index, Charge, Length, Molecular Weight, Aliphaticity, Charge Density, Boman Index, and PI. | 10 |
| Amino Acid Frequencies | The occurrence of a particular amino acid in the sequence divided by the length of the sequence. | 20 |
| Dipeptide Frequencies | The occurrence of peptides that yields two molecules of amino acid on hydrolysis. | 400 |
| Optimal Growth Temperature | The temperature at which the host organism exhibits maximum growth and reproduction. | 1 |
| **Total Number of Features** | 431 | 431 |

The Random Forest and Linear Regression models were evaluated based on the R^2 value. The Logistic Regression model was evaluated using an accuracy score. The results are shown in Table 2. Lasso Linear Regression obtained an R^2 value of 0.54 on the training set and 0.52 on the test set. Random Forest attained a R^2 value of 0.9322 on the training set and 0.624 on the test set. Logistic Regression attained an accuracy of 92.6% on the training set and 88.3% on the test set. The graphs of these results are shown in Figure 6.

**Table 2.** Results of Machine Learning models for predicting Topt.

|  | Linear Regression with Lasso Regularization | Random Forest | Logistic Regression |
| --- | --- | --- | --- |
| Training Set | $R^2 = 0.54$ | $R^2 = 0.9322$ | Accuracy = 92.6% |
| Test Set | $R^2 = 0.52$ | $R^2 = 0.624$ | Accuracy = 88.3% |

### Feature Ranking and Selection

Using Lasso Linear Regression, we ranked the enzyme features that we used as inputs for the models and algorithms. The most important 10 features are shown in Table 5. Out of 431 features, 156 features had non-zero coefficients and were kept by Lasso Linear Regression in the feature set.

### Directed Evolution for PETase Engineering

The amino acid sequence for the old original PETase enzyme is shown above in the left column of Table 4 with its Topt of 42 degrees Celsius. The Amino Acid sequence for the newly designed enzymes with 1000 and 29 iterations of directed evolution are shown in the second and third columns of Table 3. The enzyme generated with 1000 mutations has a predicted Topt of 71.38 degrees C. The enzyme generated with 29 mutations has a predicted Topt of 61.3 degrees C. Figure 5 shows the Topt of the mutant enzyme developed after 29 iterations, increasing through the iterations.

**Table 3.**
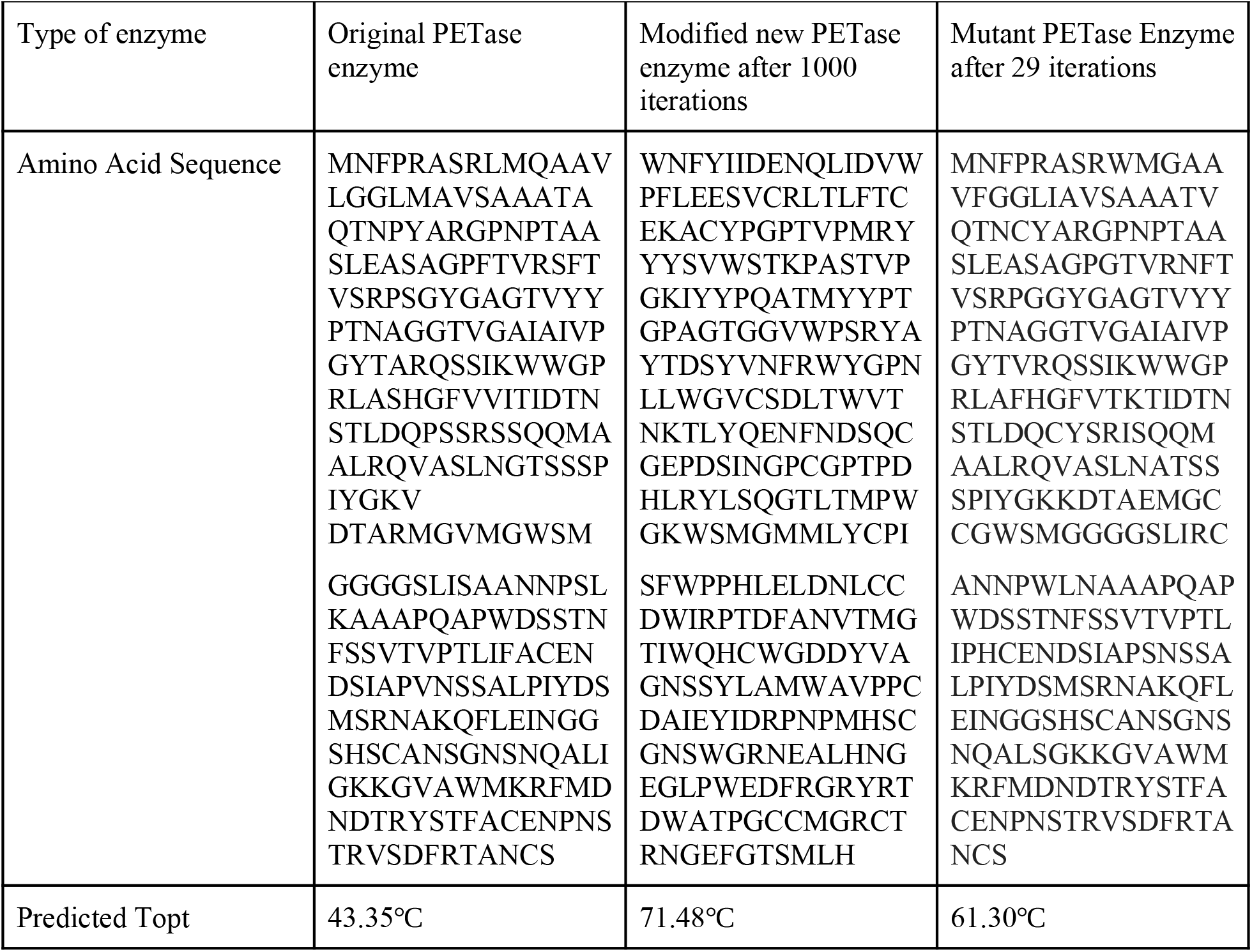
Table of comparison between original PETase enzyme and newly designed enzymes from ML-guided directed evolution.

**Table 4.**
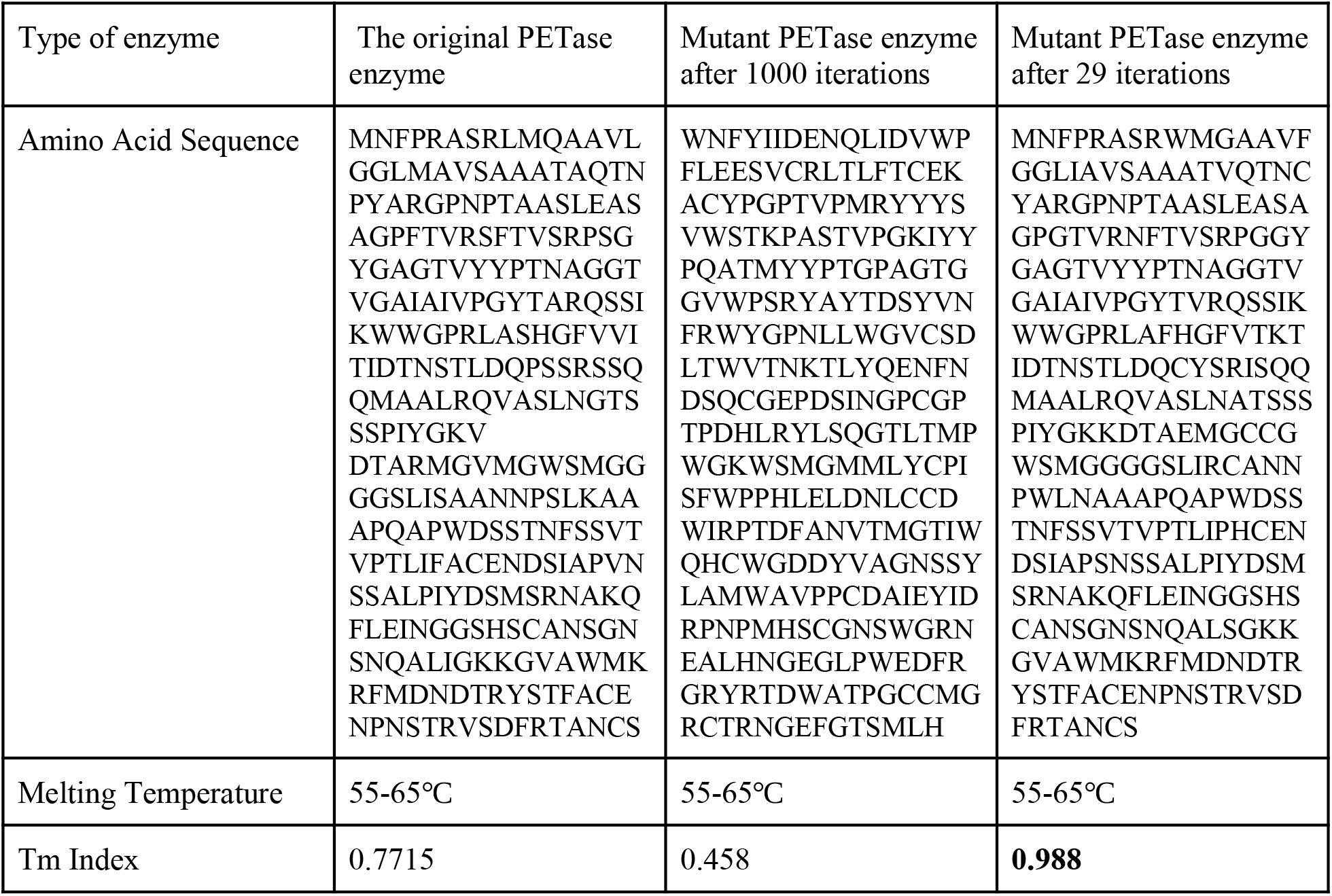
Table of predicted melting temperatures and TM Index of original PETase and mutant PETase enzymes (8).

**Figure 4:**
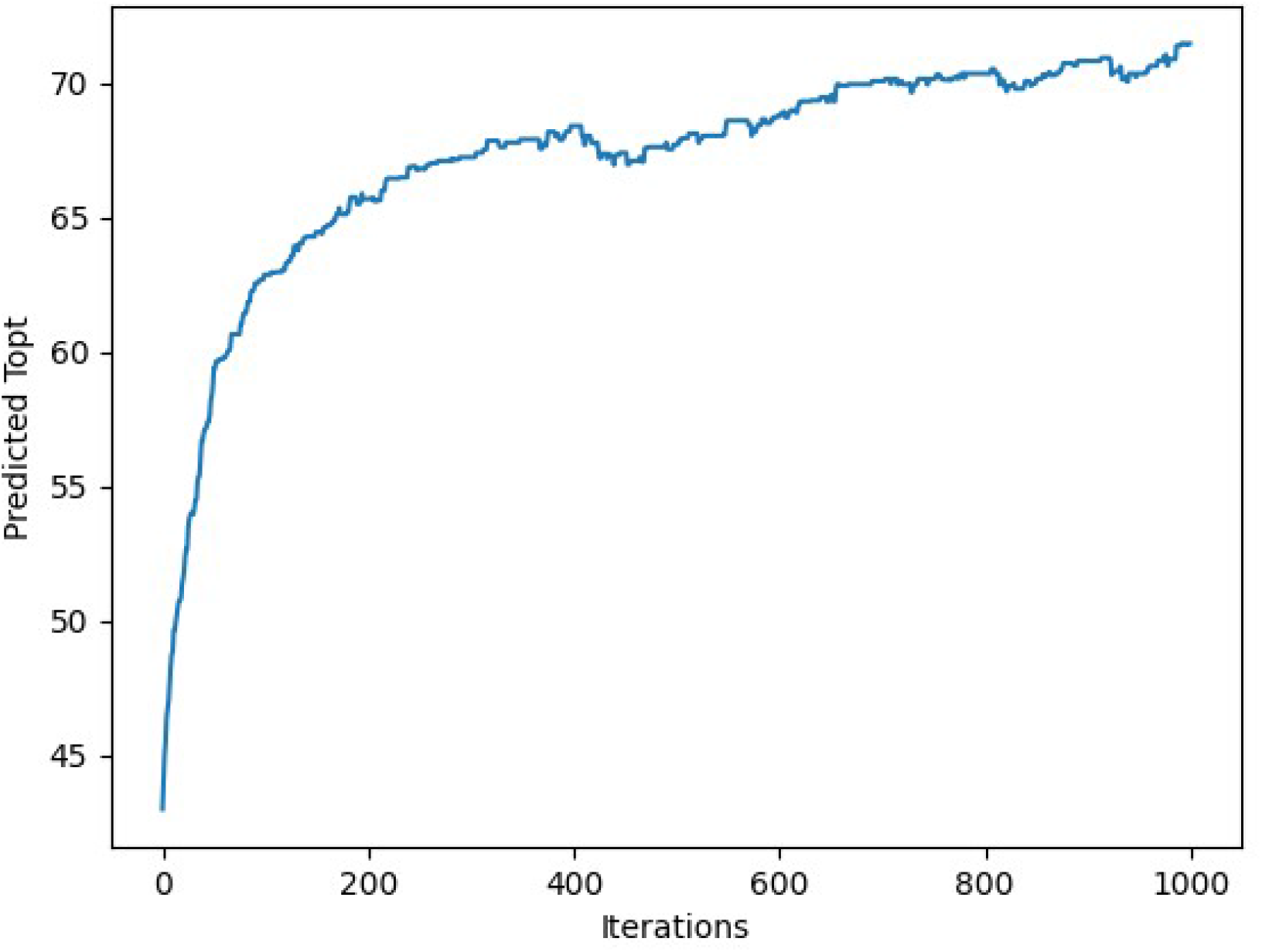
Directed Evolution graph for 1000 Iterations. This graph shows the progression of Predicted Topt we obtained over 1000 iterations of mutations and ML-guided directed evolution on the original PETase enzyme. The final Topt obtained was 71.48 degrees Celsius as seen in Table 3.

**Figure 5 :**
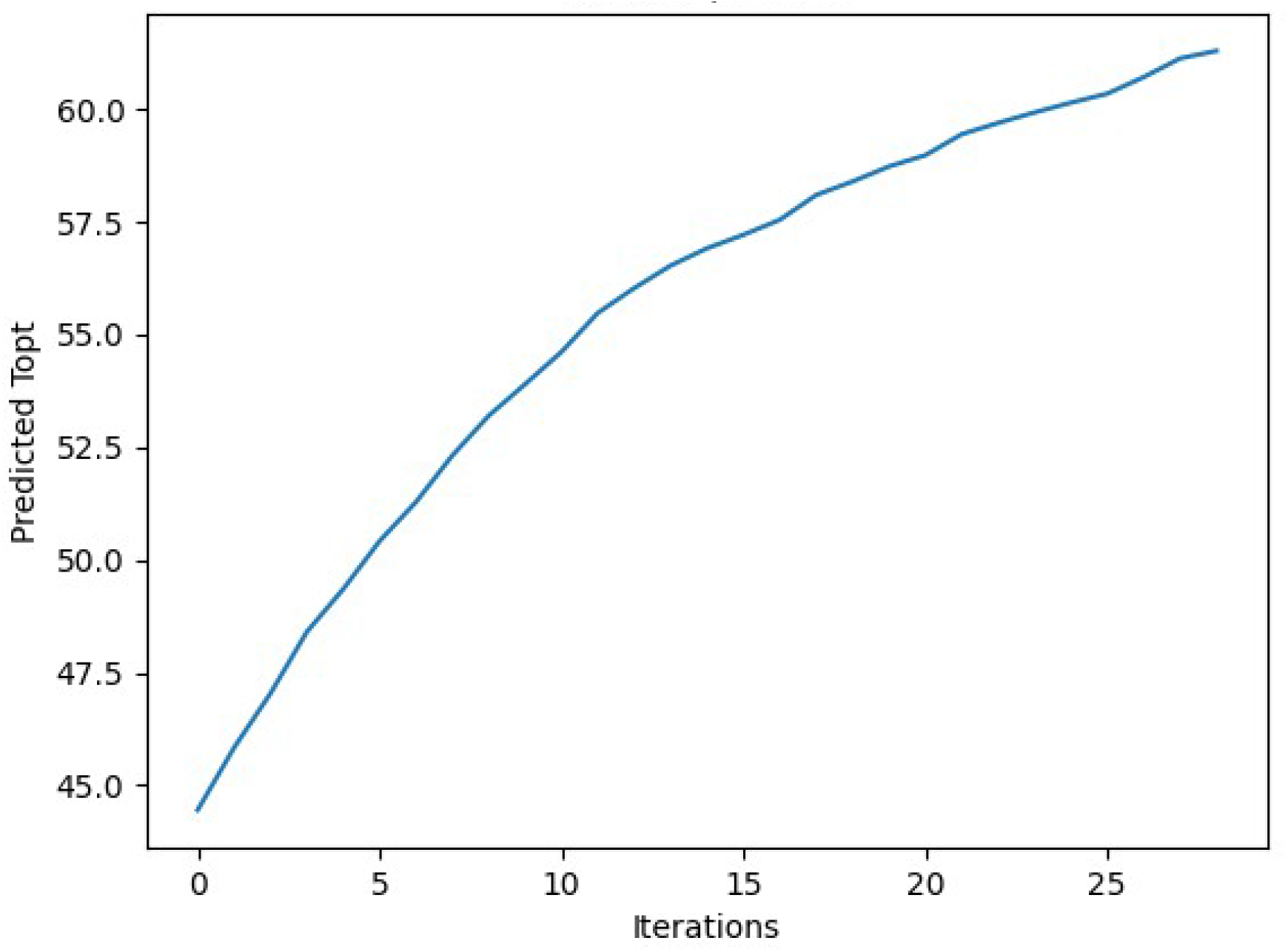
Directed Evolution over 29 iterations. This graph shows the progression of Predicted Topt we obtained over 29 iterations of mutations and ML-guided directed evolution on the original PETase enzyme. The final Topt we obtained was 61.3 degrees Celsius.

The thermostability of the enzyme was measured by the predicted melting temperatures shown in Table 4. We used the melting temperature predictor published by Ku et al (8) to predict these melting temperatures. The TM index is proportional to the actual melting temperatures of the enzymes. The TM Index > 1 implies that the True Melting temperature value of the protein may exceed 65 °C (high Tm protein), whereas a TM < 0 implies that the True Melting temperature value may be below 55 degrees C. A TM index between 0 and 1 implies that the true melting temperature is within 55 and 65 degrees Celsius. As seen in Table 4, the original PETase enzyme had a melting temperature in the range of 55 to 65 degrees C and a TM index of 0.778. The mutant PETase enzyme after 1000 iterations was also predicted as having a melting temperature in the range of 55 to 65 degrees C and achieved a TM index of 0.458. The mutant PETase enzyme after 29 iterations was also predicted as having a melting temperature in the range of 55 to 65 degrees C and received a TM index of 0.988.

## Discussion

### Machine Learning Results

Out of the three Machine learning models trained, Random Forest was the best regression model for calculating the actual Topt while Logistic Regression was the best classifier for predicting a Topt above 65 degrees C. The models were evaluated on a training and test set. A training set is the data used to build and train the model. A test set determines how well the model performs on data outside the training set. The regression models were scored based on the coefficient of determination (R^2 metric). The R2 score is a metric that is used to evaluate the performance of a regression-based machine learning model. It measures the amount of variance in the predictions explained by the dataset. The classification models were scored based on accuracy. Accuracy is the fraction of correct predictions over total predictions.

The inputs of the models and the algorithm were the enzyme features. Lasso Linear Regression was also used to rank the features by their coefficients when predicting Topt. Out of the 431 enzyme features, only 156 features had non-zero coefficients. Lasso Linear Regression kept these features in the feature set and dropped the features with the zero coefficients. The top 10 ranking features selected by Lasso Linear Regression are shown in Table 5 with their respective weights. The most important feature by far was OGT, the optimal growth temperature of the host organism of the enzyme. This makes sense because the Topt of an enzyme would naturally evolve to be around the range of the temperature where its host organism grows. For example, enzymes of thermophilic organisms would have to have a high Topt in order for the enzyme to function at the high temperatures these organisms thrive in.

**Table 5.** After feature selection by Lasso Linear Regression, only 156 features out of 431 had non-zero coefficients and were selected. The features were ranked by the absolute value of their coefficients and the top 10 features are shown in Table 5.

| Feature Name | Coefficient | Absolute Value of Coefficient |
| --- | --- | --- |
| OGT | 11.958509287023 | 11.958509287023 |
| Dipeptide Freq: YG | 1.0444261357069 | 1.0444261357069 |
| Dipeptide Freq: YN | 0.831136543632622 | 0.831136543632622 |
| Dipeptide Freq: WG | 0.733657595656585 | 0.733657595656585 |
| Amino Acid H Frequency | -0.7304045328829 | 0.7304045328829 |
| Dipeptide Freq: YL | -0.700898056815346 | 0.700898056815346 |
| Dipeptide Freq: IG | -0.6659526774771 | 0.6659526774771 |
| Amino Acid YFrequency | 0.616193055377511 | 0.616193055377511 |
| Dipeptide Freq: PG | 0.588640535816482 | 0.588640535816482 |
| Dipeptide Freq: YA | 0.558509622057296 | 0.558509622057296 |

As seen by its’ R^2 value of 0.54 on the training set and 0.52 on the test set, Linear Regression with Lasso Regularization did not work very well on both of the datasets for predicting the Topt of the enzymes. However as shown by its similar R^2 value on both the training and test set, Linear Regression generalized well. With a R^2 value of 0.9322 on the training set and 0.624, Random Forest did very well on the training set and was the best regression model for predicting Topt. Logistic Regression showed a high accuracy of 92.6 % on the training set and of 88.3% on the test set for predicting a Topt greater than or equal to 65 degrees Celsius. As compared to a previous research study that created a model to predict Topt, our algorithm performed better based on the metrics of R^2 score (9). The other model achieved a R^2 score of 0.51 on the test set using Random Forest and Deep Learning.

### Directed Evolution Results

In the directed evolution stage of the algorithm, the mutant enzyme of PETase developed as an output after 29 iterations is the better enzyme to test in the lab as it more closely resembles the original PETase enzyme as compared to the mutant enzyme developed after 1000 iterations. The mutant enzyme of PETase after 1000 iterations achieved a Topt of 71.38 degrees C whereas the mutant enzyme of PETase after 29 iterations achieved a Topt of 61.3 degrees C. As shown in Figure 4, the Topt of the enzyme increases at a very high rate for the first 200 iterations of Directed Evolution and then stabilizes and continues to rise between 65 and 70 degrees Celsius.

The mutant enzyme of PETase after 1000 iterations could be considered as a separate enzyme rather than a mutant of PETase, since its amino acid sequence differs greatly from that of the original PETase enzyme. In addition, the mutant PETase enzyme after 29 iterations achieved a much higher TM index (0.988) than the mutant PETase enzyme after 1000 iterations (0.458). This might be due to the TM predictor considering the mutant PETase enzyme after 1000 iterations to be separate from the original PETase enzyme since their amino acid sequences vary so greatly. The mutant PETase enzyme with 29 iterations more closely resembles the original PETase enzyme. These enzyme melting temperature values show that the PETase and the mutant PETase enzymes’ thermostability is between 55 and 65 degrees Celsius. This means that we can indeed optimize the optimal temperatures of PETase above 60 degrees Celsius for maximum efficiency, and the enzyme will still be stable and working at those temperatures. In addition, our 29-iteration mutated PETase enzyme has a higher TM index as compared to the original PETase enzyme, which provides external validation that our ML-guided directed evolution was successful. This also signifies that increasing the enzyme’s Topt also increased its thermostability.

### Conclusions and Future Work

In order to avoid potential sources of error, while generating mutants, the algorithm avoided mutating the substrate site of the PETase enzyme in order to ensure the enzyme would continue to function. This was implemented by preserving certain positions on the amino acid sequence and making sure the algorithm excluded those areas while mutating. Those positions corresponded to the substrate binding positions on the enzyme.

In looking towards future work on this project, we plan to synthesize and test the two mutant enzymes this algorithm produced in the lab, both after 1000 and 29 iterations. We will also test their optimal temperature for breaking down PET. We will express the enzymes into bacteria and then measure the bacteria’s efficiency in degrading PET samples of differing sizes. For a future project, I am also planning to express the best mutant enzyme in Cyanobacteria, in order to allow these photosynthetic bacteria to degrade the plastic in the sea and convert it into non-harmful products for the ocean and marine life.

## Materials and Methods

### A. Machine Learning Main Procedure and Training the Machine Learning Models

Enzyme data from the Brenda Database was used to train the machine learning models to predict the enzyme’s Topt. After obtaining the data, the algorithm split the data set into a training (90% of the data set) and independent test set (10% of the data set). The inputs for the model were the enzyme features calculated from the amino acid sequence and are listed in Table 1. These inputs include amino acid frequency, dipeptide frequency, and optimal growth temperature (OGT). Optimal growth temperature is defined as the temperature at which the host organism of the enzyme has maximum growth and reproduction. OGT was listed in the BRENDA database. The remaining features were calculated from the amino acid sequences through the modLAMP python library(12). Before performing a cleanup of the BRENDA data set, there were 2745 enzyme amino acid sequences comprising the dataset. During the cleanup process, we used the Pandas library to drop duplicate rows, amino acid sequences with a length less than or equal to 7, and enzymes with a Topt equal to or lower than 0 degrees Celsius. The final training data consisted of 2643 enzyme amino acid sequences listed with the experimental Topts of each enzyme.

Three Machine learning models were trained to predict Topt: Random Forest, Linear Regression and Logistic regression. Classification Models such as logistic regression are concerned with predicting a discrete label, such as Topt >= 65 degrees C. Regression models such as Linear Regression and Random Forest models fundamentally predict a continuous quantity such as the actual Topt value for the enzymes. The three Machine Learning models were implemented using the scikit python library (11).

In the training stage, first Linear Regression was trained to have a high R^2 value (correlation value) between the Predicted Topt and the actual Topt of the enzymes. However in ordinary least squares regression, the R^2 values were only 0.607 for the training set and 0.33 for the test set. This showed that Linear regression was overfitting and not generalizing well to unseen data from the test set. Thus, we had to regularize Linear regression by using the lasso method. Lasso Regression is linear regression with added l1 regularization. This regularization adds a penalty to a model if the model is overfitting, meaning if it is setting the value of its weights too high. Here,Lasso regularization set the weights of the enzyme features that were unimportant to Linear Regression to 0. Now after implementing Lasso, Linear regression obtained an R^2 value of 0.54 on the training set and 0.52 on the test set.

Then a Random Forest regression model was trained which performed much better on minimizing error between the predicted and actual Topt. Random Forest models operate by assembling different decision trees, then taking the mean of those decision trees and setting that as the predicted output. Random Forest attained a R^2 value of 0.9322 on the training set and 0.624 on the test set.

Logistic Regression was the classifier that was trained and it was measured by its accuracy in predicting whether an enzyme’s Topt was above 65 degrees Celsius. Logistic Regression attained an accuracy of 92.6% on the training set and 88.3% on the test set. The graphs of these results are shown in Figure 6.

**Figure 6:**
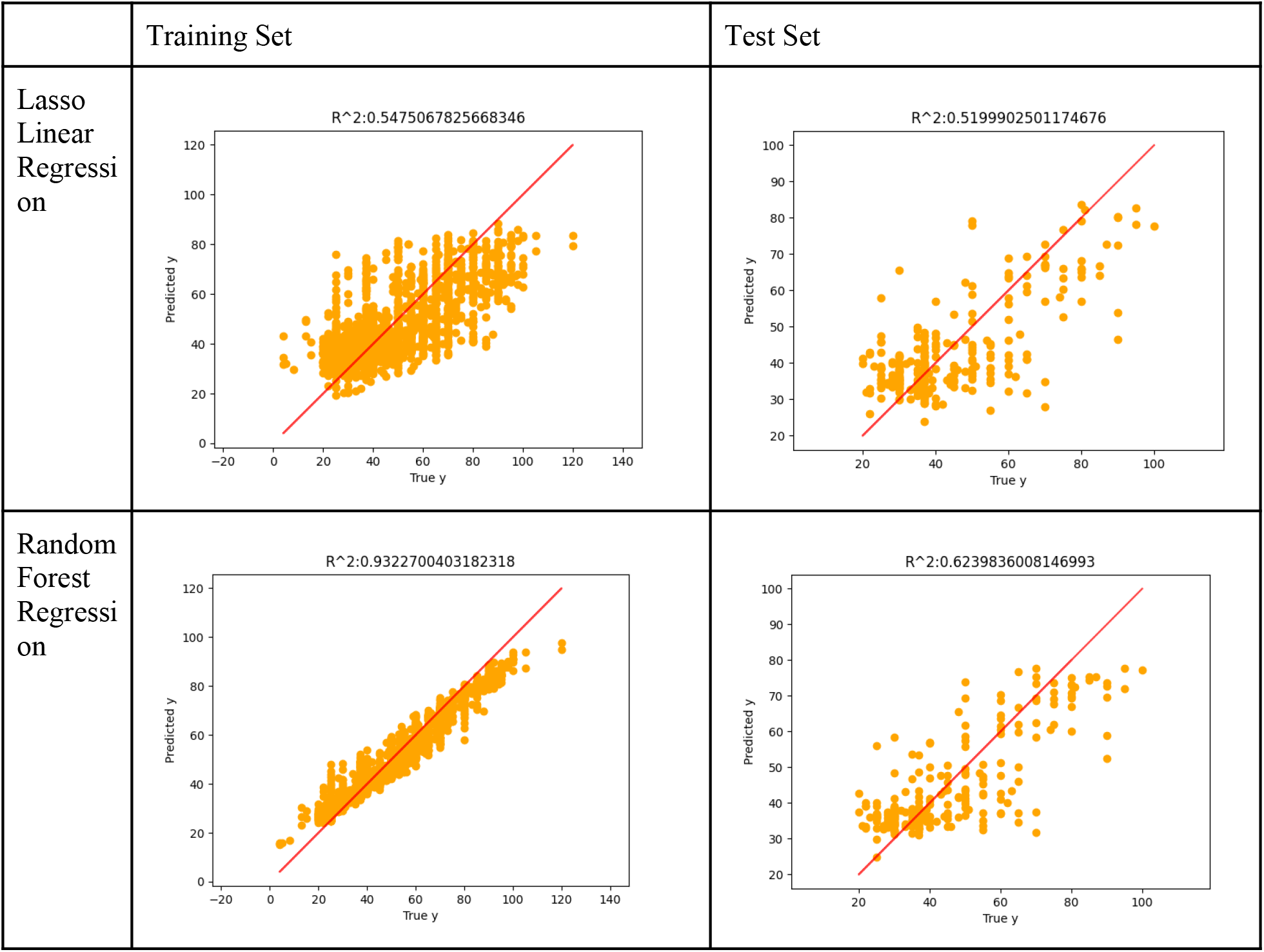
Graphs for Machine Learning training stage results: These graphs show the performance of the regression machine learning models, Lasso Linear Regression and Random Forest Regression, on the training and test set. The correlation between the predicted Topt values and actual Topt values is shown through the R^2^ coefficient.

### B. Directed Evolution Procedure

The ML-guided directed evolution algorithm generated a thousand mutants of PETase on every iteration and screened them using Random Forest Regression. The enzyme with the highest Topt was then selected and mutated in the next iteration of the algorithm. This process is known as machine learning (ML) guided *in silico* directed evolution and it was used in order to steer the mutants of PETase towards higher thermostability. In this type of directed evolution, machine learning is used to score and evaluate different possible mutations of enzymes. Based on the machine learning scores, the algorithm then selects the best mutant and uses it as a starting point again.

First, in our ML-guided directed evolution approach, the algorithm randomly mutated the PETase enzyme at random positions, excluding the substrate site at positions where the PET molecule binds to the enzyme according to the Uniprot database (13). The algorithm did not mutate the binding site of the enzyme because this might decrease enzymatic activity. A thousand mutants were generated in this way. Second, based on those mutations, the algorithm used Random Forest to score the corresponding mutants based on their predicted Topt values. The best scoring mutant was then selected based on its optimal temperature. The algorithm then repeated this mutation and selection process with the top mutant from the previous iteration now acting as the main enzyme. Summed up, after performing directed evolution on the mutants of PETase, the algorithm selected the best scoring mutant and performed random mutations on it at random positions, excluding the substrate site. This process was repeated for 1000 iterations and yielded an enzyme with 71.38 degrees C Topt. However, 1000 iterations would lead to approximately 1000 mutations in the original enzyme, which would create an enzyme vastly different from the original PETase. The original PETase enzyme has only 290 amino acids. Thus, this process was then also repeated for 29 iterations in order to yield a mutant enzyme more similar to PETase rather than a completely new enzyme. After 29 iterations, ML-guided directed evolution produced an enzyme with a predicted Topt of 61.3 degrees C.

### C. External Validation of Mutant PETase Enzyme

The enzyme’s melting temperature ranges and TM index values were generated by the online melting temperature predictor by Ku *et al* (8). The TM values are shown in Table 4. These melting temperatures further validate that the mutant enzymes produced by the algorithm are stable and can function at their optimal temperatures without breaking down. These values show that the PETase and the mutant PETase enzymes’ thermostability is between 55 and 65 degrees Celsius. The TM index for the mutated PETase enzyme is larger than that of the original PETase enzyme, signifying that increasing the enzyme’s Topt also increased the enzyme’s thermostability.

